# How adaptive plasticity evolves when selected against

**DOI:** 10.1101/339622

**Authors:** Alfredo Rago, Kostas Kouvaris, Tobias Uller, Richard Watson

## Abstract

Adaptive plasticity allows organisms to cope with environmental change, thereby increasing the population’s long-term fitness. However, individual selection can only compare the fitness of individuals within each generation: if the environment changes more slowly than the generation time (i.e., a coarse-grained environment) a population will not experience selection for plasticity even if it is adaptive in the long-term. How does adaptive plasticity then evolve? One explanation is that, if competing alleles conferring different degrees of plasticity persist across multiple environments, natural selection between lineages carrying those alleles could select for adaptive plasticity (lineage selection).

We show that adaptive plasticity can evolve even in the absence of such lineage selection. Instead, we propose that adaptive plasticity in coarse-grained environments evolves as a by-product of inefficient short-term natural selection. In our simulations, populations that can efficiently respond to selective pressures follow short-term, local, optima and have lower long-term fitness. Conversely, populations that accumulate limited genetic change within each environment evolve long-term adaptive plasticity even when plasticity incurs short-term costs. These results remain qualitatively similar regardless of whether we decrease the efficiency of natural selection by increasing the rate of environmental change or decreasing mutation rate, demonstrating that both factors act via the same mechanism. We demonstrate how this mechanism can be understood through the concept of learning rate.

Our work shows how plastic responses that are costly in the short term, yet adaptive in the long term, can evolve as a by-product of inefficient short-term selection, without selection for plasticity at either the individual or lineage level.

## Introduction

Organisms that live in variable environments are often subject to opposing selective pressures, either temporal or spatial, such that intermediate generalist phenotypes have decreased fitness across all environments. Rather than evolving a generalist phenotype, populations can keep adapting to each environmental condition as they encounter them, a process known as adaptive tracking [1, 2]. Populations that evolve via adaptive tracking avoid the trade-offs paid by generalist phenotypes, but need time to adapt to each new environment. Consequently, the population experiences reduced fitness after each environmental change. Both populations that evolve a generalist phenotype and those that evolve by adaptive tracking thus have reduced fitness in the long term. By contrast, adaptive phenotypic plasticity allows individuals to maintain an adaptive fit between phenotype and environment: plastic individuals produce only high fitness phenotypes by responding appropriately to environmental cues. Populations evolving adaptive plasticity thus avoid both the fitness loss arising from trade-offs of generalist phenotypes and the fitness loss that tracking populations suffer after environmental change. Within this framework, the question of whether plasticity evolves can be interpreted as the comparison between the long-term average fitness of populations which evolve plastic responses, evolve generalist phenotypes or evolve via tracking [3, 4]. As such, a considerable amount of effort has been invested in characterizing the conditions that determine the fitness of plastic rather than non-plastic solutions, and to document if plasticity itself incurs a fitness cost [5–7].

While adaptive plasticity is common in nature and demonstrably superior to non-plastic solutions for a wide range of conditions, the process by which it evolves remains a matter of debate. The standard assumption that natural selection favours the best available option is problematic, since natural selection only discriminates between phenotypes that are expressed. Natural selection is thus unable to detect that a plastic organism is adapted to more environments than a non-plastic one unless individuals encounter multiple environments within their life spans, a condition known as environmental fine-grain [8]. Even when individuals experience more than one environment per lifetime, plastic individuals may still be limited to a single phenotype if plastic responses are irreversible [9–11], too slow (e.g [12]) or too costly (e.g. [5]) relative to the fitness advantage of producing the right phenotype for the local conditions [2, 7].

Each of those cases creates an evolutionary trap: adaptive plasticity provides the best long-term solution, but natural selection favours non-plastic phenotypes in each short-term environment. In other words, experiencing one environment per lifetime (environmental coarse grain) does not allow individual selection for plasticity, so that if plastic responses incur any cost compared to non-plastic phenotypes they will be selected against in the short-term. Nevertheless, many examples of adaptive plasticity have been reported even when organisms experience only coarse-grained environments. Examples of adaptive responses to coarse-grained environments include the production of resistance and dispersal forms [13, 14] and seasonal morphs of short-lived species [9, 15].

How can we explain the process by which costly adaptive plasticity evolves in such coarse-grained environments? While individual-level selection does not favour plasticity in coarse-grained environments, alleles that determine an organism’s plasticity are transmitted between generations, and their fixation or loss will depend on their fitness across the set of environments they encounter [16, 17]. Natural selection may therefore discriminate between plastic and non-plastic alleles if both are maintained long enough to be selected across multiple environments, even if each individual organism experiences only a single environment. Plastic adaptations to coarse-grained environments could therefore evolve via genetic lineage selection [4, 18, 19].

A key requirement for the evolution of adaptive plasticity via lineage selection is the availability and persistence of standing genetic variation on plastic responses (e.g. [4, 17, 19]). This implies that plasticity will not evolve in populations that are small or under strong selection, since these conditions remove the genetic variation lineage selection requires to operate (e.g. [20]). Because small population size and strong selection are representative for populations experiencing rapid environmental change, evolution of plasticity appears unlikely to play a role in evolutionary rescue or successful colonization [21, 22]. The evolution of costly adaptive plasticity will only be possible if genetic diversity is available, but high genetic diversity will also cause rapid removal of costly plastic variants in favour of non-plastic short-term solutions, so that costly adaptive plasticity should only evolve as an intermediate step towards non-plastic solutions.

We apply a core concept of learning theory - learning rate - to propose an alternative mechanism for the evolution and maintenance of costly adaptive plasticity without lineage selection. In machine learning, learning rate measures the amount of change a system accumulates with each example shown. Existing literature demonstrates that the process of learning by trial and error is mechanistically analogous to evolution by natural selection [23]. In the context of adaptation, genetic learning rate measures the ability of a population to change in response to new environments by accumulating adaptive mutations. More specifically, we can define genetic learning rates as the amount of genetic change fixed by a population in each new environment. Genetic learning rate (henceforth just learning rate) depends both on the ability to generate variation (mutation rate and effect size, population size) and to fix particular variants (strength of selection). Since both the processes that produce and fix variants require time to operate, increasing the time spent in each environment will allow populations to accumulate more adaptive change. Thus, the more generations a population spends in a single environment the higher its learning rate will be.

As we show in our simulations, populations initially produce phenotypes matching their current environment by accumulating both mutations that change the mean phenotypic value and mutations that change its plasticity. Populations with high learning rates find optimal short-term phenotypes and remove costly plasticity before each new environmental shift: Efficient selection in each short-term environment prevents the evolution of costly long-term adaptive plasticity. Populations with low learning rates cannot reach short-term optima before the next environmental shift, and pass on to the next environment all genetic changes which brought them closer to the previous phenotypic optimum, whether or not these genetic changes cause phenotypes to be plastic. Short-term selection in the new environment thus starts from a population which already accumulated adaptively plastic changes, so that the overall plastic responses can be further refined over time. In evolutionary terms, low learning rates maintain directional selection for plastic development with the end result of directing evolution towards the production of long-term adaptive plastic responses.

Unlike the lineage selection explanation, the learning theory explanation does not require the prolonged co-existence of alleles with different effects on plasticity: adaptive plastic responses will evolve even in populations which exhibit only a single reaction norm at any given time. Rather, learning theory only requires that the population accumulates limited genetic change per environment, so that the average genotype retains some of the adaptive plasticity accumulated in past environments. Learning theory thus predicts that, as long as natural selection is inefficient, long-term adaptive plasticity should evolve even in the extreme case when only one lineage is present in the population (strong selection weak mutation) and plasticity is selected against in the short-term.

In this paper, we provide a first exploration of the evolution of adaptive plasticity from a learning theory perspective. To do so, we employ a classic linear reaction norm model [24, 25] to simulate the evolution of costly adaptive plasticity in temporally coarse-grained scenarios. This allows us to contrast the predictions made by learning theory and lineage selection regarding when and how plasticity should evolve.

First, we demonstrate that plasticity can evolve in coarse-grained environments, showing that individual-level selection for plasticity is not necessary to evolve adaptive plasticity. Second, we demonstrate that adaptive plasticity evolves in coarse-grained environments even in the absence of multiple lineages, counter to the predictions of lineage selection. Third, we show that limiting mutation rates biases populations towards adaptive plasticity rather than adaptive tracking.

These results are consistent with the predictions of learning theory and reveal that long-term adaptations can evolve even when short-term fitness selects against them, as long as natural selection is inefficient.

## Results and Discussion

### Simulation Set-up

We simulate a population that experiences temporal environmental heterogeneity. Each individual receives information from the environment and develops into an adult phenotype, upon which selection can act. We follow standard approaches for the evolution of plasticity [18, 26, 27] and model development as a linear reaction norm, whose intercept *a* represents the genetic trait value and slope *b* the degree of plasticity (see Reaction Norm Model). The developed phenotype *P* is thus

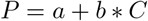

where *C* is the univariate environmental cue.

We model a heterogeneous varying environment with 10 environmental states, so that each environmental state, *E_i_* produces a single, unique value of the cue *C^E_i_^* and requires a single specific univariate phenotype *P^E_i_^*. We model the matching between cues and trait optima as a linear function (see Environmental Variability). This implies that a linear reaction norm with appropriate slope and intercept can achieve perfect fit for all environments in our set. We assume non-overlapping generations of individuals with a constant fixed lifespan. This assumption allows us to control the granularity of environmental variability with a single parameter, *K*. If *K* ≥ 1 the environment changes every *K* generations, indicating coarse-grained (*K* = 1) or slow coarse-grained (*K* > 1) environmental variability. If instead *K* < 1 the population encounters on average 1/*K* environments per generation, indicating fine-grained environmental variability.

We evaluate the fitness of each individual based on the distance of its developed phenotype from the optimal target phenotype in the current environment. In case the individuals experience more than one environment, we calculate their fitness as the mean match between the developed phenotypes and the selective environments experienced. We further impose a fitness penalty proportional to the individual’s responsiveness to its environment (reaction norm slope *b*, see above). This cost of plasticity ensures that plastic individuals will have lower fitness than non-plastic ones regardless of their phenotypes, and effectively represents a trade-off incurred by plastic organisms (see Evaluation of Fitness). Organisms reproduce asexually with a probability proportional to their relative fitness within the population (see Evolutionary Process). Every individual inherits the same slope and intercept as their parents, which are then mutated by adding a random value selected from a normal distribution with mean 0 and standard deviation equal to the mutation size (0.01 unless otherwise specified). Thus, both intercept and slope mutate every generation (effective mutation rate = 1), but most mutations have small effects.

Unless otherwise stated, we set a population of 1000 individuals and choose a selection coefficient ω of 0.2. In addition, we set the associated cost of plasticity, λ, to be 0. 1. We analytically tested all parameter combinations used in our simulations and confirmed that the fitness losses caused by adaptive tracking exceed those caused by the cost of plasticity for all of them (see S1 Appendix). This allows us to rule out the explanation that plasticity evolves as the result of a change in its long-term fitness compared to adaptive tracking.

**Fig 1.**
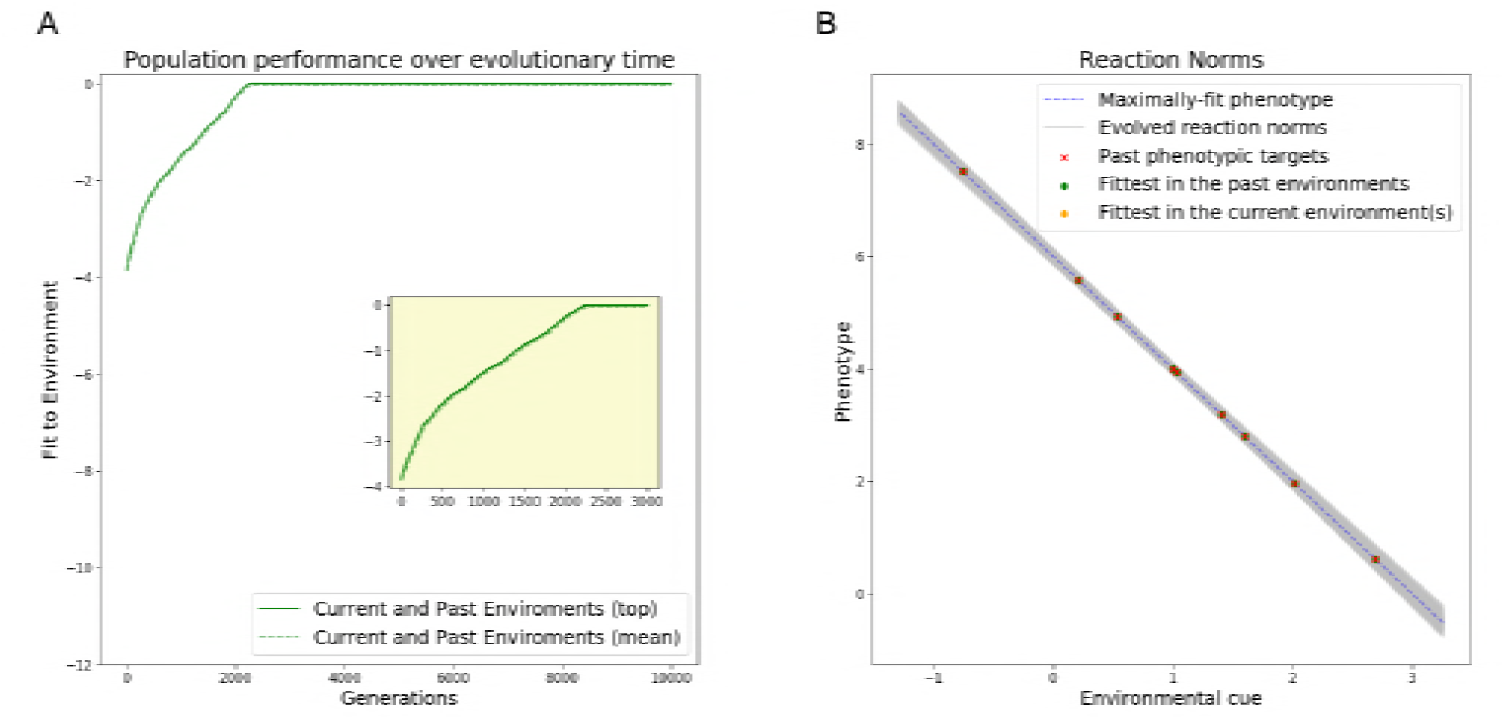
Evolution of reaction norms in fine-grained environments. (A) Average (dashed line) and best (solid line) performance of the population over time, relative to the optimal adaptive reaction norm, see Evaluation of Reaction Norms (B) Evolved reaction norms (grey lines) compared to optimal reaction norm (dashed line) at the end of the evolutionary period. Crosses indicate optima corresponding to environmental values used in the simulation. Dots indicate the phenotypes expressed for environmental values used in the simulation. The population evolves optimal adaptive plasticity

### Individual-level selection is not necessary for the evolution of adaptive plasticity

In this section, we compare the evolution of plasticity in fine-grained environments, which allow individual-level selection for plasticity, with coarse-grained ones, which do not. We initially assess the evolution of phenotypic plasticity when individuals encounter multiple environmental states per life-time (i.e., a fine-grained environment; here 10, *K* = 0.1). We further assume that the phenotype can change during individuals’ lifespan (reversible plasticity), and this change is both immediate and incurs in no fitness costs.

In fine grained environments, the evolved reaction norms converge the optimal intercept and slope in less than 3000 generations (Fig 1A, inset). This means that individuals produce trait values that perfectly match the optimal trait value of all environmental states they encountered during their lifetime, as we can see from the fact that the distance between realised and optimal phenotypes decreases to zero for all environments in our set (Fig 1A). We find minimal residual genetic variation on both the slope and intercept terms of the reaction norm (Fig 1B). This is reflected in the limited differences between the reaction norms of top and mean performing individuals (Fig 1A). Note that the reaction of the average (yellow dots) and best individual (green dots) are perfectly aligned and match the optimal reaction norm (red crosses).

We contrast the previous fine-grained scenario with a slow coarse-grained environment in which the local conditions change every 4000 generations on average (*K* = 4000). As such, each individual experiences only one environment, and environmental change between generations is also slow. In this coarse-grained environment, the population fails to evolve adaptive long-term plasticity (Fig 2). After each environmental change we observe a drop in short-term fitness, followed by a distinctive two-step pattern in their adaptive paths. During the first phase, organisms evolve towards the new target phenotype, as indicated by the steep increase in short-term fitness (Fig 2A, inset, green line). Crucially, the increase in short-term fitness in this phase is accompanied by a corresponding increase in long-term fitness (Fig 2A, blue line), which indicates evolution of adaptive plasticity. Mutations which increase plasticity can be selected for during this phase if they cause the production of fitter phenotypes, offsetting their fitness cost (see S1 Appendix). After organisms are able to produce phenotypes which match the short-term phenotypic optima, we observe a decrease in their long-term fitness (Fig 2A, blue curve). This indicates that the same organisms would no longer be able to produce adaptive phenotypes when exposed to past environments, consistent with a decrease in costly adaptive plasticity. During this phase plasticity is directly selected against in order to decrease its fitness costs.

**Fig 2.**
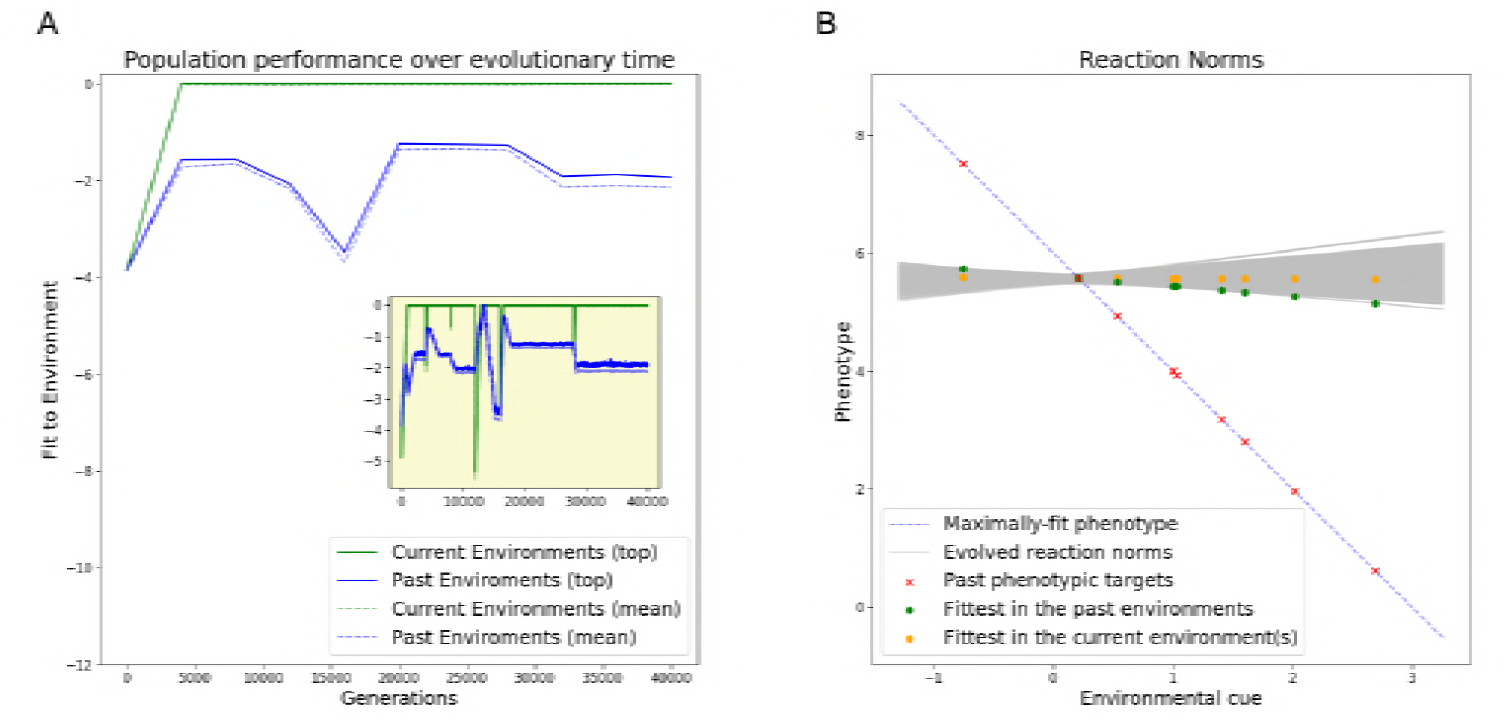
Evolution of reaction norms in slow coarse-grained environments. (A) Relative performance (see Evaluation of Reaction Norms) in the local (green lines) and global (blue lines) environments. Dashed lines indicate average population performance, solid ones best performance. (B) Evolved reaction norms (grey lines) compared to optimal reaction norm (dashed line) at the end of the evolutionary period. Crosses indicate optima corresponding to environmental values used in the simulation. Dots indicate the phenotypes expressed for those values. The population re-adapts to the local environment after each environmental change (adaptive tracking)

In other words, the population reaches the optimal phenotype using a combination of slope and intercept (**phenotypic adaptation**) and then minimizes the slope (**plasticity minimization**). From a fitness perspective, selection during the phenotypic adaptation phase increases fitness by producing the local target phenotype, whereas selection in the plasticity minimization phase increases fitness by removing costly plasticity. It is worth noting that these two phases match those described in the analogous model presented in [17]. After the plasticity minimization phase we still observe some genetic variation in reaction norm slope (grey lines in Fig 2B), but the average slope is 0: adaptive plastic responses are approximately as likely as maladaptive ones. Populations evolving under slow, coarse-grained environments thus fail to evolve adaptive plasticity and instead re-adapt upon each environmental change, consistently with adaptive tracking.

Next, we test whether direct selection for plasticity is required for its evolution. To do so, we set the environment to change every generation (*K* = 1), which is the fastest rate we can set under a coarse-grain scenario: every individual experiences only a single environment, but every generation experiences a different one. Since each individual only experiences one environment, we can rule out direct selection for adaptive plasticity. Furthermore, costly plasticity is selected against within each short-term environment.

**Fig 3.**
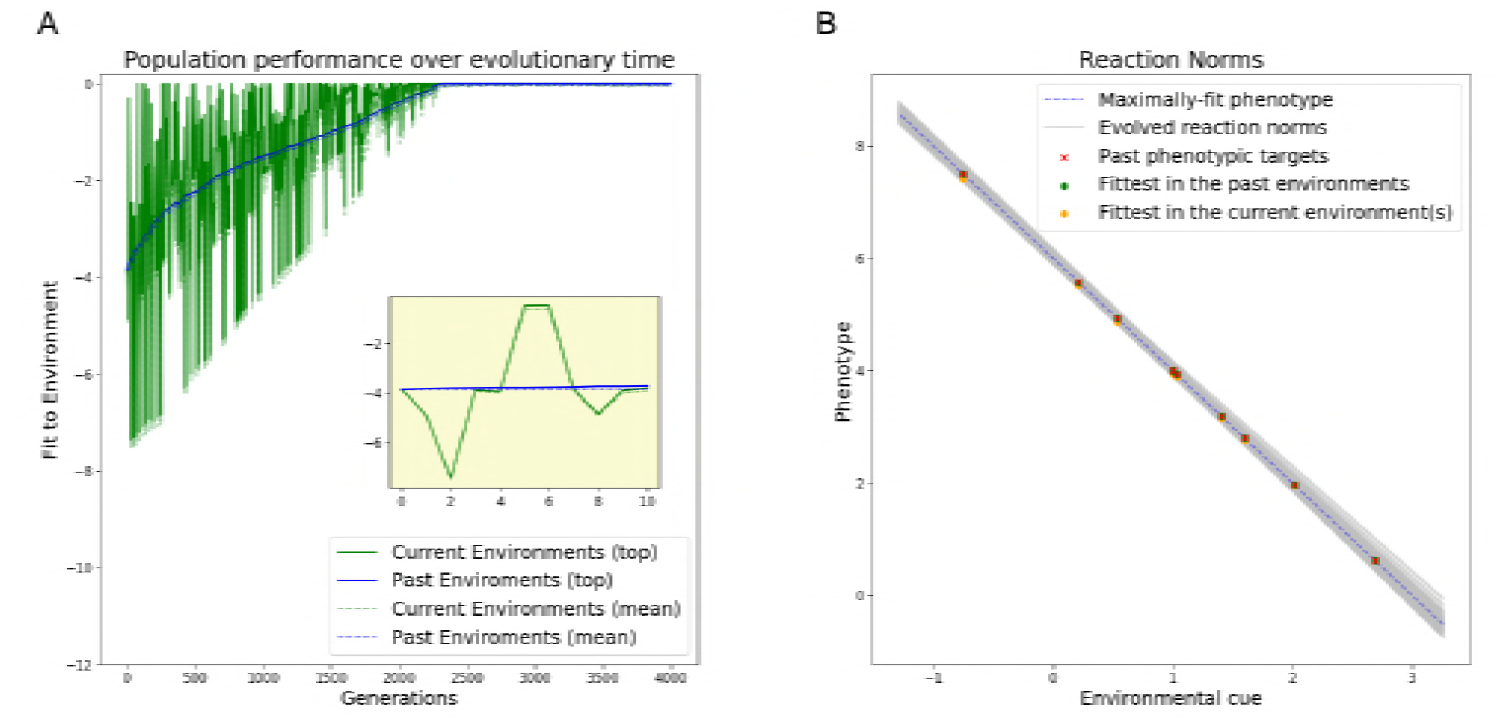
Evolution of reaction norms in fast coarse-grained environments. (A) Relative performance (see Evaluation of Reaction Norms) in the local (green lines) and global (blue lines) environments. Dashed lines indicate average population performance, solid ones best performance. (B) Evolved reaction norms (grey lines) compared to optimal reaction norm (dashed line) at the end of the evolutionary period. Crosses indicate optima corresponding to environmental values used in the simulation. Dots indicate the phenotypes expressed for those values. The population evolves optimal adaptive plasticity.

In this fast coarse-grained environment, populations evolved adaptive plasticity (Fig 3). We observe that the goodness of fit to current and past environments decreased to zero, indicating optimal fit to all environments within the range experienced (Fig 3A). In addition, we observe less residual genetic variation compared to the case of slow coarse-grained environmental variability (Fig 3B). This is also indicated by the narrow gap between the top and the mean performance curve in Fig 3A.

Looking at the evolutionary trajectory of the population, we can see that while fitness to the current environment (i.e., short-term environment; green line) fluctuates, fitness to the whole environment set (i.e., long-term environment; blue line) gradually increases over time. Moreover, we see no gap between performance in the short-term environment and in the long-term environment. This indicates that the population does not evolve short-term fit phenotypes at the expense of long-term performance, but rather directly accumulates responses that are adaptive in the long-term environment. These results demonstrate that populations evolving in fast-changing environments produce adaptive plastic responses even when plasticity is costly and environmental change only occurs between generations.

At this stage, we have merely confirmed well-known results (e.g., [17]). We now consider two explanations for the evolution of adaptive plasticity in coarse-grained environments. According to a lineage selection model, faster environmental change will increase the odds that each allele is tested in more than one environment. Adaptive plasticity can evolve since plastic alleles have greater mean fitness than non-plastic, short-term adaptive, alleles when compared across multiple environments. The learning theory model instead predicts that decreasing the number of generations in each environment will decrease the genetic change accumulated within each environment (i.e., the learning rate), ensuring that the changes accumulated during the phenotypic adaptation phase are not lost because of short-term optimization. While both mechanisms cause a shift from short to long-term adaptation, each has distinct requirements: lineage selection relies on the transmission of genetic variation in order to compare the fitness of multiple alleles; learning theory requires that populations accumulate little genetic change in each environment. In the next two sections, we make use of this key difference to determine which of the two processes can better explain the evolution of plasticity in coarse environments.

### Lineage selection is not necessary for the evolution of adaptive plasticity

In order to test the need for lineage selection, we repeat the scenarios for the evolution of plasticity in fine-grained (*K* = 0.1), coarse-grained (*K* = 1) and slow coarse-grained (*K* = 40000) environments enforcing strong selection and weak mutation (SSWM). Under SSWM, the speed at which mutations arise is much slower compared to the speed at which they are fixed or lost, driving standing genetic variation to zero. Comparing the fitness of alleles across different environments is therefore impossible. We model SSWM using a hill-climber algorithm: each evolutionary step produces only one mutation. If the new mutation is fitter than the previous one it is fixed, otherwise it is lost (see Hill-climbing Model). SSWM leads to a constant effective population size of 1 and makes lineage selection impossible. Therefore, if the lineage selection hypothesis is correct, we expect that adaptive plasticity will fail to evolve in all coarse-grained environments. To rule out that the potential failure to evolve plasticity is due to insufficient time, we verify the results under an extended simulation time of 2 * 10^7^ generations.

Contrary to the predictions of the lineage selection explanation, we find that the results from the above simulations are qualitatively and quantitatively similar to those obtained using a population size of 1000, despite the SSWM selection regime (Fig 4). That is, populations fail to evolve plasticity when environments change every 40000 generations (Fig 4A), and succeed in doing so when provided with either fine environmental grain (Fig 4B) or a rapid coarse-grained (i.e., trans-generational) change (Fig 4C).

The evolutionary trajectory of populations under SSWM also remains remarkably similar to that of populations with standing genetic variation (compare Fig 4 with Fig 1, Fig 2 and Fig 3). Populations evolving in fine-grained and fast coarse-grained environments both show a gradual increase in long-term fitness, which remains comparable to short-term fitness. This indicates that they evolve by adapting to the long-term environment rather than to the short-term environments. Populations in slow coarse-grained environments instead perform consistently better in short-term, current, environments compared to the long-term one, showing the repeated evolution of short-term adaptive phenotypes, or adaptive tracking. Their evolutionary trajectory also displays the same two-step cycle after each environmental change: fitness increase in both short and long-term environments (phenotypic adaptation) followed by fitness decrease in the long-term environment only (plasticity minimization) (Fig 4A).

Taken together, these findings demonstrate that both the final results and the evolutionary trajectories of our simulations are largely unaffected by the lack of standing genetic variation. Since standing genetic variation is required for adaptation via lineage selection, these results falsify the hypothesis that plasticity needs to evolve by averaging the fitness benefits of alternative variants across multiple environments. In the next section, we make predictions based on the learning theory explanation and try to falsify them.

**Fig 4.**
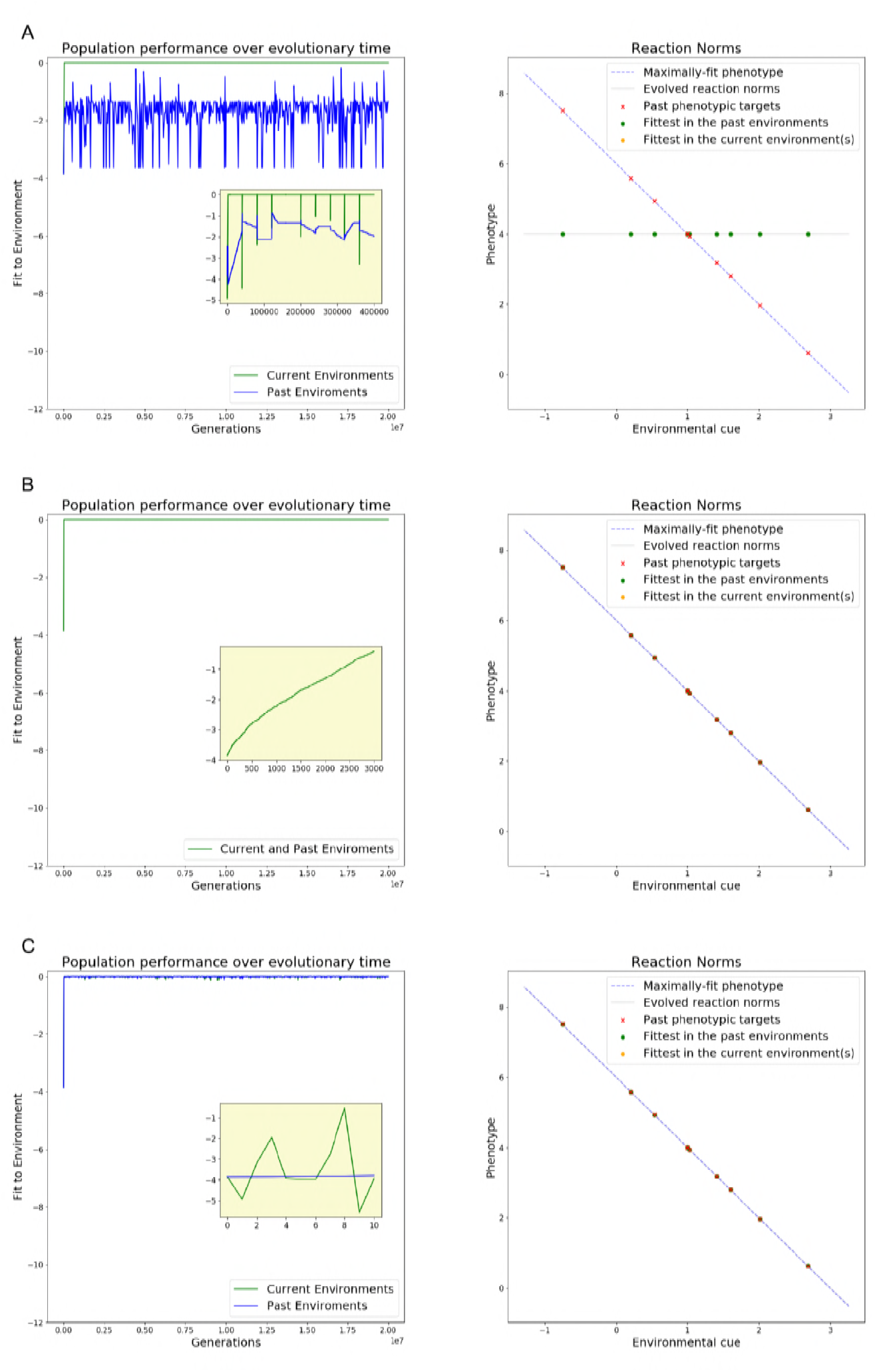
Evolution of reaction norms under Strong Selection Weak Mutation. Panels to the left show population performance (see section Evaluation of Reaction Norms) over time in the local (green line) and global (blue line) environment. Panels to the right show the evolved reaction norm (solid line) compared to optimal reaction norm (dashed line) at the end of the evolutionary period. Crosses indicate optima corresponding to environmental values used in the simulation. Dots indicate the phenotypes expressed for those values. (A) Slow coarse-grained environments (K=40000) (B) Fine-grained environments (K=0.1) (C) Fast coarse-grained environments (K=1). Performance over time and evolved reaction norms are identical to weak selection scenarios

### Low mutation rates are analogous to fast environmental change

Since we define learning rates in biological systems as the amount of genetic change accumulated in each new environment, they can be affected by several parameters other than rate of environmental change. Population size, mutation size and mutation frequency will all increase the amount of genetic change produced within each environment and thus increase the population’s learning rates. Stronger selective pressure will speed up the fixation of beneficial variants, and therefore also increase learning rates. If the learning rate explanation for the evolution of adaptive plasticity in coarse-grained environments is correct, these factors should be interchangeable with the rate of environmental change.

In this section, we evaluate the learning theory explanation by testing the specific prediction that adaptively plastic responses can evolve even when environmental changes are slow, provided that mutation sizes are sufficiently small (and hence learning rate is low). In order to test this prediction, we return to the case of slow coarse-grained environments (environments change every 40000 generations) with a population size of 1000 individuals. As shown above, adaptive plasticity fails to evolve under these conditions. Learning theory explains this failure with the high learning rates in this population. Rather than decreasing the learning rate by decreasing the number of generation spent in each environment, we lower the standard deviation of mutation sizes from 10^−2^ to 10^−4^.

As we can see in Fig 5B, the population eventually evolves an optimally adaptive plastic reaction norm, with negligible amounts of variation around both slope and intercept. Their evolutionary trajectories (Fig 5A) are also qualitatively similar to those of populations evolving in fast, coarse-grained environments. In both scenarios, fitness in the short-term environment (green) fluctuates around average fitness in the long-term environment (blue), indicating that the populations are not evolving phenotypes that increase short-term fitness at the expense of long-term adaptation. The steady increase in average fitness instead indicates the evolution and retention of more general, plastic solutions.

While the two trajectories are similar in shape, the population experiencing slower environmental changes and smaller mutation rates takes a significantly longer amount of time to reach optimal plasticity. An increase in the number of generations required to find solutions is a known consequence of lower learning rates. Intuitively, we can explain the longer time required to adapt as a consequence of the slower rate at which variants become available.

While lineage selection is technically viable in this simulation, decreasing mutation sizes would also decrease the amount of available genetic variation, making it even less effective. Therefore, it is unlikely that lineage selection causes the recovery of plasticity in mutation limited populations. Nevertheless, in order to ensure that the results we observe are not caused by lineage selection, we run a simulation with *K* = 40000 and *σ_μ_* = 10^−5^ using a hill-climber to model SSWM. The results are both qualitatively and quantitatively similar to those obtained in the previous simulation (see Fig 6). Since we are unable to falsify the learning rate hypothesis, we conclude that it provides the best explanation for evolution of costly adaptive plasticity under coarse-grain.

## Conclusion

The evolution of costly adaptive plasticity has often been framed as a necessity caused by environmental change outpacing the ability of natural selection to generate new adaptations [2, 3, 28, 29], but the means by which organisms achieve plasticity in these conditions have seldom been clarified.

We demonstrate that neither individual nor lineage-level selection for adaptive plasticity are necessary for it to evolve. Rather, the speed of adaptation relative to environmental change (modelled as learning rates) is by itself a causal factor in the evolution of adaptive long-term plastic responses. High learning rates allow local optimization of phenotypes in each environment, at the expense of more general solutions. Low learning rates instead make it impossible for phenotypes to chase short-term optima, yet allow individuals to reach long-term optimal plasticity despite the presence of short-term trade-offs.

**Fig 5.**
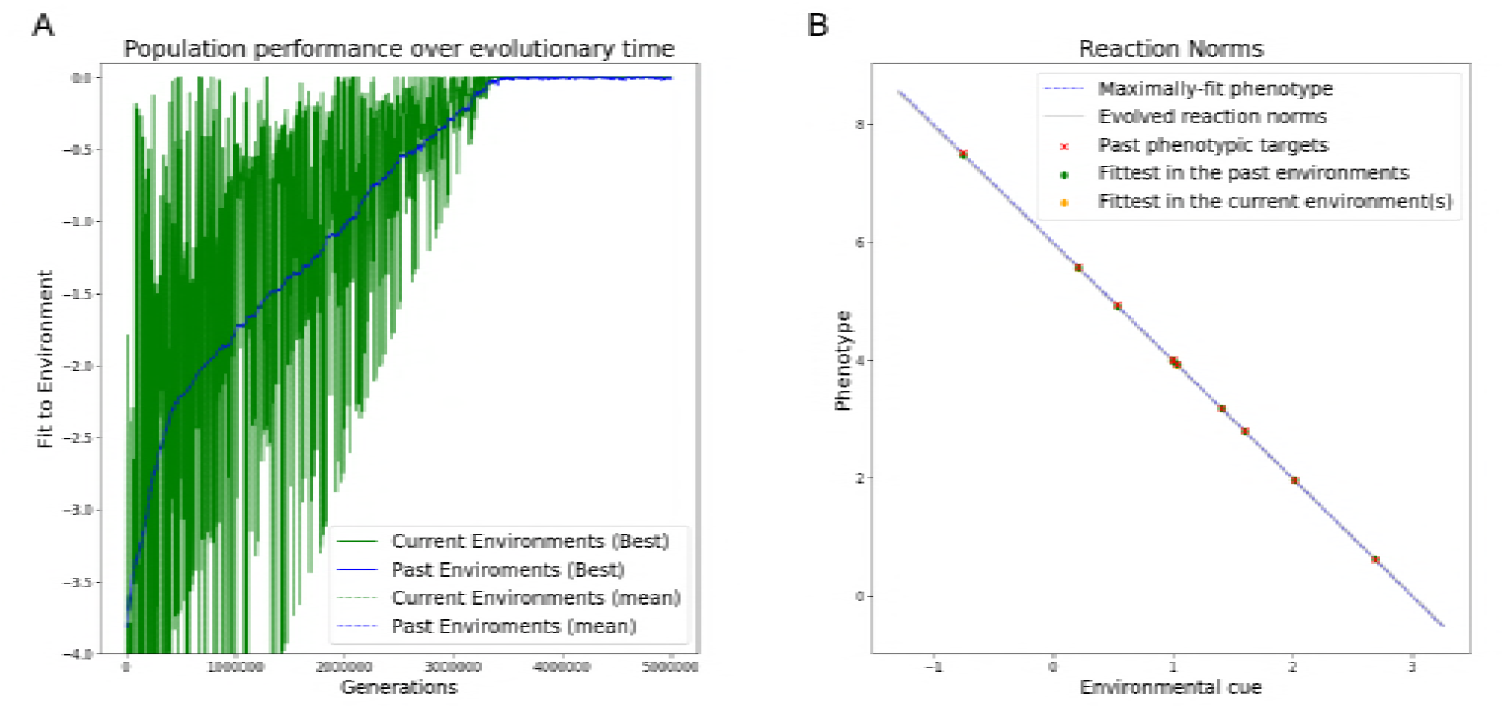
Evolution of reaction norms in slow coarse-grained environments with low mutation rates. (A) Relative performance (see Evaluation of Reaction Norms) in the local (green lines) and global (blue lines) environments. Dashed lines indicate average population performance, solid ones best performance. (B) Evolved reaction norms (grey lines) compared to optimal reaction norm (dashed line) at the end of the evolutionary period. Crosses indicate optima corresponding to environmental values used in the simulation. Dots indicate the phenotypes expressed for those values. The population slowly evolves optimal adaptive plasticity.

**Fig 6.**
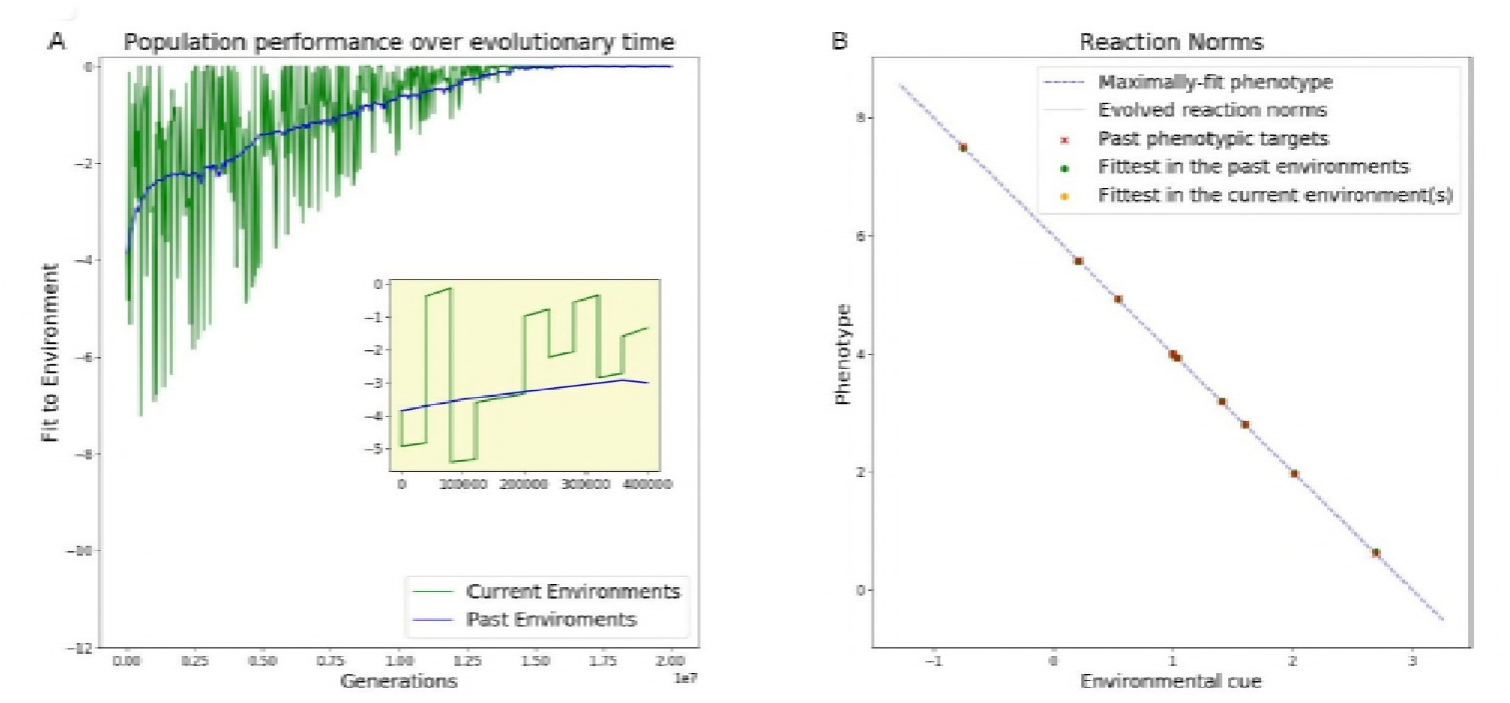
Evolution of reaction norms in slow coarse-grained environments with low mutation rates under SSWM. (A) Relative performance (see Evaluation of Reaction Norms) in the local (green lines) and global (blue lines) environments. (B) Evolved reaction norms (grey lines) compared to optimal reaction norm (dashed line) at the end of the evolutionary period. Crosses indicate optima corresponding to environmental values used in the simulation. Dots indicate the phenotypes expressed for those values. The population slowly evolves optimal adaptive plasticity.

Although plasticity evolves when natural selection is least effective, plasticity maximizes long-term population fitness in all our simulations. In contrast, adaptive tracking produces higher fitness phenotypes in the short term, but shows lower fitness in the long term due to the fitness losses that follow each environmental change (see S1 Appendix).

While low learning rates are necessary to evolve general solutions in the presence of trade-offs in performance, none of the factors that affect learning rates is necessary by itself. This is because learning rate is a composite measure, so any given factor may be offset by the others. We demonstrate this by showing that low mutation rate is sufficient to evolve costly adaptive plasticity even in slow, coarse-grained environments. Increasing population size and selection strength will instead decrease the odds of evolving costly adaptive plasticity, as both factors increase learning rates. As a consequence, even populations with no measurable genetic variation in plasticity could evolve adaptive plastic responses as long as (1) new genetic variation can be produced over time and (2) environmental change is faster than adaptation.

This observation reverses the suggested causal link between plasticity and the rate of genetic evolution. Current theory proposes that plastic individuals experience weaker selection because they are able to cope with a wider range of environments [4]. Because of the reduced selective pressure, the amount of genetic change that accumulates in the population (learning rate) is also reduced. We instead suggest a low learning rate itself may skew populations towards evolving more general solutions, such as short-term costly but long-term optimal adaptive plasticity.

Since low learning rates promote the evolution of adaptive plastic responses by reducing the relative importance of minimizing plasticity costs, they will be irrelevant to the evolution of inexpensive plastic responses. If there are no costs of plasticity, every combination of slope and intercept that generates the optimal short-term phenotype will be fitness equivalent within each environment. This implies that adaptive plasticity will be selected for while the population moves towards the local phenotypic optimum and randomly drift after the optimal phenotype has been reached. The population will thus inevitably find the long-term optimum, and learning rates will only determine the speed of the adaptation and drift processes.

Learning rates are likewise irrelevant for the evolution of costly adaptive plasticity in fine-grained environments, which are sufficient (but not necessary) for the evolution of adaptive plasticity across all our simulations. Fine-grained environments allow natural selection to directly compare the fitness of phenotypes across multiple environments at the individual-level within each generation, so that adaptive plasticity is optimal even in the short-term. Direct selection for plasticity is unsurprisingly sufficient to ensure the evolution of adaptive plasticity, so that learning rates can only determine the speed of selective process rather than its outcome.

Our simulations consider the specific case of maintenance costs for plasticity. That is, we assume that plasticity directly decreases fitness, regardless of whether it is expressed. However, several alternative scenarios can create mathematically equivalent trade-offs between short and long-term selection. A well-studied example is that of inaccurate cues, either due to imperfect perception or noise in the cues themselves [3, 21, 30]. Alternatively, the target phenotypes may not perfectly match with the best possible reaction norm. This scenario can happen for any reaction norm which is selected on a set of environments larger than its degrees of freedom (3 in the case of linear reaction norms) [31] or if there are limits to the maximum amount of plastic changes that an organism can evolve [26, 32–34]. In all of the above mentioned cases, optimal long-term plasticity would cause loss of fitness in the short term and be consequently selected against. Learning rates will thus be relevant for the evolution of plastic responses across all of them.

In our simulations, mutations that lead to adaptive plasticity are selected since they increase phenotypic fitness within the short-term environment, and adaptive long-term plasticity evolves as a result of direct selection for fitter phenotypes in the short term. This is in contrast with lineage selection models, in which adaptive plasticity evolves because of its long-term benefits, but is (at best) selectively neutral within each environment. Since the evolution of plasticity in our model is driven by a direct (rather than lineage) selection process, we predict it to be both faster and more robust to the presence of short-term trade-offs. Similar dynamics apply to the evolution of modularity as a by-product of short-term phenotypic selection, and are proven to be scalable to arbitrarily complex systems in which the long-term optimal solutions incur short-term fitness costs [35].

From a learning theory perspective, low learning rates cause the evolution of adaptive plasticity because they constrain populations to evolve new adaptive solutions starting from previous adaptations rather than ‘from scratch’. As a result, evolved reaction norms do more than just ‘remember’ the specific instances of cue to phenotype matchings experienced during selection: they capture the logic that matches cues and phenotypes. In learning theory terms, organisms learn the regularities of the (evolutionary) problem, a process also known as generalization [36]. Therefore, as long as regularities remain the same, each individual will be able to produce adaptive phenotypes even in environments it has never experienced in its evolutionary history (extrapolation). Several studies show that systems which learn a problem’s regularities are also able to quickly adapt to new problems which share a similar logic, increasing their evolvability [37, 38]. Our demonstration that organisms can learn regularities between environments even when they do not experience them within their lifetimes opens up the possibility that evolved plastic responses may allow organisms to both anticipate future environments and enable them to more rapidly evolve novel adaptive solutions. This demonstrates that past evolution can shape evolutionary trajectories by biasing the phenotypic variants that are exposed to selection [23].

In summary, we use a simple reaction norm model to demonstrate that costly adaptive plasticity can evolve even when natural selection is unable to compare competing alleles over multiple environments (i.e., lineage selection). A learning theory framework helps us interpret this finding: Populations evolving in coarse-grained environments can evolve adaptive plasticity if the amount of adaptive change accumulated per environment - the learning rate - is low. Populations with high learning rates evolve via repeated short-term adaptation even if this pattern is maladaptive in the long term. Low learning rates facilitate long-term adaptation over short-term adaptation, favouring adaptive plasticity even in the presence of short-term functional trade-offs. Thus, long-term adaptive plasticity can evolve even when it is not selected for at either the individual nor lineage level. Whether a population evolves phenotypes that optimize fitness in the short or long term instead depends on the amount of adaptive changes it accumulates within each environment.

## 1 Methods

### 1.1 Environmental Variability

We model a heterogeneous global environment in which each population is exposed to local environments with different phenotypic optima, each characterized by reliable cues. We assume that the lifespan of the individuals is fixed and the same for all. As a result, environmental grain is solely determined by the parameter *K. K* < 1 indicates fine-grained environmental variability, where the population encounters an average of 1/*K* environments per generation. On the other hand, *K* >= 1 indicates coarse-grained (*K* = 1) or slow coarse-grained (*K* > 1) environmental variability where the population encounters a new environment every *K* generations on average. We choose small *K* values compared with the total number of generations in our simulations so that each population is able to evolve for multiple environmental cycles.

For plasticity to evolve, the environment needs to fulfil two roles: determining the selective conditions (selective role) and providing information about those conditions (constructive role) [39]. We simulate the selective role by assigning each local environmental state a target single trait optimum *ϕ*, represented by a single real number. We simulate the constructive role by assigning each target optima an environmental cue represented by a real number *C*, which varies between 0 and 1. For simplicity, we consider a linear relationship between phenotypic targets and environmental cues, so that *ϕ* = *g*(*e*) = *g*_1_ * *e* + *g*_0_. Hence, the targets are directly proportional to the respective cue. We choose *g*_1_ = −2 and *g*_0_ = 6. This ensures that the relationship between selective environment and cues remains constant across environmental states.

Our simulations were designed with temporal variation in mind, but the conclusions should be applicable to spatial variation as well. In fact, the environmental fluctuations described within our model match those experienced by a population in which all individuals migrate after fitness evaluation and before reproduction, or in which all propagules are dispersed to the same new environment. In this scenario environmental change rates are effectively interchangeable with migration rates, with other findings remaining unchanged.

### 1.2 Reaction Norm Model

We model plastic responses using a univariate linear reaction norm model [40]. A reaction norm can be defined as the set of phenotypes that would be expressed if the given individual would be exposed to the respective set of environments. Since we consider univariate and linear reaction norms, we can describe the development of an organism’s phenotype as *P* = *a* + *b* * *C*. Each organism’s genotype can thus be described by the factors *a* and *b*. Of those, a determines the organism’s breeding value and b the direction and magnitude of its plasticity.

### 1.3 Evolutionary Process

We model the evolution of a population of asexual individuals as follows. First, we select a parent using a fitness proportional criterion [41, 42]. Each individual can be selected with a probability of 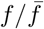, where 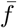 corresponds to mean fitness in the current population and *f* to the parent’s own fitness (see section 1.4 for details on how we calculate *f*). Then, we generate a new individual with the same genotype (reaction norm intercept *a* and slope *b*) as the parent. Finally, we independently mutate both the offspring’s intercept and slope by adding a random value sampled from a normal distribution with mean *μ* = 0 and standard deviation equal to mutation size (*σ_μ_* = 0.01 unless otherwise specified). We repeat this process until we generate a number of offspring equal to the set population size. The parameters *a* and b are initialized at zero.

### 1.4 Evaluation of Fitness

Following previous work [31, 35, 37], we define an organism’s overall fitness *f* in terms of a benefit-minus-cost function, which allows us to consider both positive (benefits) and negative (costs) contributions to its fitness. The benefit of a given genotype, *b^E_i_^*, for each environment, *E_i_*, is determined based on how close the developed adult phenotype, *P^a^*, is to the target phenotype, *P^E_i_^*, of the given selective environment, *E_i_*. Since we deal with an univariate phenotype, we can calculate this amount as

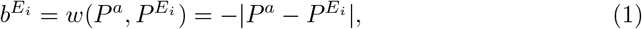

where |*| corresponds to the absolute distance between the two phenotypes. Note that the selective advantage of respective genotypes is solely determined by its immediate fitness benefits on the currently encountered selective environment(s). We consider that individuals experience a distribution of selective environments during their lifetime with occurring probabilities, *q*^*E*_1_^, *q*^*E*_2_^,…, *q^E_N_^*. Each environment contributes to the selection process in proportion to its occurrence [43]. The overall fitness benefits of an individual over all experienced environments in its lifetime, *b^E^* is determined by the arithmetic mean of the fitness benefits in each environment, *b^E_i_^*, weighted by the occurrence, *q^E_i_^*, of each environment:

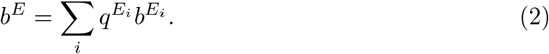

In cases of coarse-grained environmental variability, where each individual encounters a single environment in its lifespan, *q^E_i_^* = 1 for the respective environment, *i* = *j*, and *q^E_i_^* = 0 for *i* ≠ *j*. On the other hand, in cases of fine-grained environmental variability, we assume a uniform distribution of environments experienced during individual’s lifespan, that is, *q^E_i_^* = 1/*K*. The cost represents how maintaining plasticity reduces the organism’s fitness. Unlike the benefit, the cost of plasticity is a property of the genotype and does not change in different environments. Thus, we can calculate the overall performance, *d*, of a genotype over a range of selective environments as

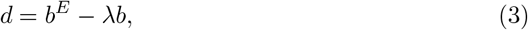

where parameter λ indicates how steeply fitness decreases in proportion to the reaction norm slope *b*. The final fitness score is calculated with the following formula:

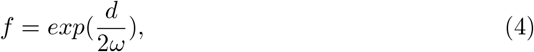

which penalizes lower performances exponentially and re-scales them to a 0-1 range. ω is a scaling factor on the relation between *f* and *d*. Lower *ω* values cause greater loss of fitness per loss of performance, and correspond to steeper selection gradients. We choose *ω* = 0.2, which corresponds to a scenario of strong selection (see [37]).

### 1.5 Evaluation of Reaction Norms

We evaluate the adaptive potential of the population due to plasticity by estimating how close the reaction norm of each individual in the population is to the (theoretical) optimal reaction norm. The optimal reaction norm here corresponds the function that given any environmental cue, *C^E_i_^*, produces the appropriate target phenotype, *P^E_i_^*, which best matches the local selective environment, *E_i_* (Evaluation of Fitness). We evaluate the performance of reaction norms based on how well they fit the optimal reaction norm. The goodness of fit, *Perf_D_* of a given reaction norm, *D*, is estimated as a function of the phenotypic trait values in each of the past selective environments (here 10), *E_i_*, that quadratically decreases with the distance from each phenotypic optimum, *p^E_i_^*:

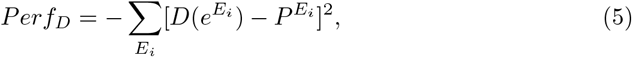

The evaluation of goodness of fit is performed for each individual at the end of each environmental period. We report the average and best performance in the population.

### 1.6 Hill-climbing Model

A hill-climbing evolutionary model simulates a scenario of strong selection and weak mutation, where each new mutation is either fixed or lost before a new one can arise. Therefore, the entire population shares the same values of *a* and *b*. Each evolutionary step introduces a single mutant genotype with parameters *a*′ and *b*′ equal to *a* and *b* plus a random value sampled from a normal distribution with mean 0 and standard deviation equal to mutation size. We develop both the reference and mutant phenotypes *P* and *P*′ (section 1.2) and compare their fitness values *f* and *f*′ (section 1.4). If *f*′ > *f*, the mutation is beneficial and therefore adopted so that *a*_*t*+1_ = *a*′ and *b*_*t*+1_ = *b*′. Otherwise, the mutation is deleterious and *a* and *b* remain unchanged.

## Supporting information

**S1 Appendix. Fitness Cost of Plasticity and Tracking** Numerical calculations of the expected long-term fitness costs of adaptively plastic and adaptive tracking populations.

## References

1. Simons AM. Modes of response to environmental change and the elusive empirical evidence for bet hedging. Proceedings of the Royal Society B: Biological Sciences. 2011;278(1712):1601–1609. doi:10.1098/rspb.2011.0176.

2. Botero Ca, Weissing FJ, Wright J, Rubenstein DR. Evolutionary tipping points in the capacity to adapt to environmental change. Proceedings of the National Academy of Sciences. 2015;112(1):184–189. doi:10.1073/pnas.1408589111.

3. Moran NA. The Evolutionary Maintenance of Alternative Phenotypes. The American Naturalist. 1992;139(5):971–989. doi:10.1086/285369.

4. Ghalambor CK, McKay JK, Carroll SP, Reznick DN. Adaptive versus non-adaptive phenotypic plasticity and the potential for contemporary adaptation in new environments. Functional Ecology. 2007;21(3):394–407. doi:10.1111/j.1365-2435.2007.01283.x.

5. Hoverman JT, Relyea RA. How flexible is phenotypic plasticity? Developmental windows for trait induction and reversal. Ecology. 2007;88(3):693–705. doi:10.1890/05-1697.

6. Van Buskirk J, Steiner UK. The fitness costs of developmental canalization and plasticity. Journal of Evolutionary Biology. 2009;22(4):852–860. doi:10.1111/j.1420-9101.2009.01685.x.

7. Auld JR, Agrawal AA, Relyea RA. Re-evaluating the costs and limits of adaptive phenotypic plasticity. Proceedings of the Royal Society B: Biological Sciences. 2010;277(1681):503–511. doi:10.1098/rspb.2009.1355.

8. Levins R. Evolution in changing environments: some theoretical explorations. 2. Princeton University Press; 1968.

9. Rountree DB, Nijhout HF. Hormonal control of a seasonal polyphenism in Precis coenia (Lepidoptera: Nymphalidae). Journal of Insect Physiology. 1995;41(11):987–992. doi:10.1016/0022-1910(95)00046-W.

10. Moczek AP. Pupal remodeling and the evolution and development of alternative male morphologies in horned beetles. BMC evolutionary biology. 2007;7:151. doi:10.1186/1471-2148-7-151.

11. Michie LJ, Masson A, Ware RL, Jiggins FM. Seasonal phenotypic plasticity: Wild ladybirds are darker at cold temperatures. Evolutionary Ecology. 2011;25(6):1259–1268. doi:10.1007/s10682-011-9476-8.

12. Kaji T, Palmer AR. How reversible is development? Contrast between developmentally plastic gain and loss of segments in barnacle feeding legs. Evolution. 2017;71(3):756–765. doi:10.1111/evo.13152.

13. van Gestel J, Weissing FJ. Regulatory mechanisms link phenotypic plasticity to evolvability. Scientific Reports. 2016;6(1):24524. doi:10.1038/srep24524.

14. Gerber N, Kokko H, Ebert D, Booksmythe I. Daphnia invest in sexual reproduction when its relative costs are reduced. Proceedings of the Royal Society B: Biological Sciences. 2018;285:20172176. doi:10.1098/rspb.2017.2176.

15. Brakefield PM, Reitsma N. Phenotypic plasticity, seasonal climate and the population biology of Bicyclus butterflies (Satyridae) in Malawi. Ecological Entomology. 1991;16(3):291–303. doi:10.1111/j.1365-2311.1991.tb00220.x.

16. Via S, Gomulkiewicz R, De Jong G, Scheiner SM, Schlichting CD, Van Tienderen PH. Adaptive phenotypic plasticity: consensus and controversy. Trends in Ecology and Evolution. 1995;10(5):212–217. doi:10.1016/S0169-5347(00)89061-8.

17. Lande R. Adaptation to an extraordinary environment by evolution of phenotypic plasticity and genetic assimilation. Journal of Evolutionary Biology. 2009;22(7):1435–1446. doi:10.1111/j.1420-9101.2009.01754.x.

18. Gomulkiewicz R, Kirkpatrick M. Quantitative Genetics and the Evolution of Reaction Norms. Evolution. 1992;46(2):390–411.

19. Siljestam M, Östman. The combined effects of temporal autocorrelation and the costs of plasticity on the evolution of plasticity. Journal of Evolutionary Biology. 2017;30(7):1361–1371. doi:10.1111/jeb.13114.

20. Volis S, Ormanbekova D, Yermekbayev K, Song M, Shulgina I. Introduction beyond a species range: A relationship between population origin, adaptive potential and plant performance. Heredity. 2014;113(3):268–276. doi:10.1038/hdy.2014.25.

21. Reed TE, Waples RS, Schindler DE, Hard JJ, Kinnison MT. Phenotypic plasticity and population viability: the importance of environmental predictability. Proceedings of the Royal Society B: Biological Sciences. 2010;277(1699):3391–3400. doi:10.1098/rspb.2010.0771.

22. Chevin LM, Lande R. When do adaptive plasticity and genetic evolution prevent extinction of a density regulated population? Evolution. 2010;64(4):1143–1150. doi:10.1111/j.1558-5646.2009.00875.x.

23. Watson RA, Szathmáry E. How Can Evolution Learn? Trends in Ecology & Evolution. 2016;31(2):147–157. doi:10.1016/j.tree.2015.11.009.

24. Schmalhauzen II. Factors of Evolution: The Theory of Stabilizing Selection. Blakiston; 1949.

25. de Jong G. Phenotypic Plasticity as a Product of Selection in a Variable Environment. The American Naturalist. 1995;145(4):493–512. doi:10.1086/285752.

26. Via S, Lande R, May N. Genotype-Environment Interaction and the Evolution of Phenotypic Plasticity. Evolution. 1985;39(3):505–522.

27. de Jong G. Quantitative Genetics of reaction norms. Journal of Evolutionary Biology. 1990;3(5-6):447–468. doi:10.1046/j.1420-9101.1990.3050447.x.

28. Fischer B, Taborsky B, Kokko H. How to balance the offspring quality-quantity tradeoff when environmental cues are unreliable. Oikos. 2011;120(2):258–270. doi:10.1111/j.1600-0706.2010.18642.x.

29. Gomez-Mestre I, Jovani R. A heuristic model on the role of plasticity in adaptive evolution: plasticity increases adaptation, population viability and genetic variation. Proceedings of the Royal Society B: Biological Sciences. 2013;280(1771):20131869–20131869. doi:10.1098/rspb.2013.1869.

30. Panchanathan K, Frankenhuis WE. The evolution of sensitive periods in a model of incremental development. Proceedings of the Royal Society B: Biological Sciences. 2016;283(1823):20152439. doi:10.1098/rspb.2015.2439.

31. De Jong G. Evolution of phenotypic plasticity: patters of plasticity and the emergence of ecotypes. New Phytologist. 2005;166(2003):101–118. doi:10.1111/j.1469-8137.2005.01322.x.

32. de Jong G. Quantitative Genetics of reaction norms. Journal of Evolutionary Biology. 1990;3(5-6):447–468. doi:10.1046/j.1420-9101.1990.3050447.x.

33. Auld JR, Agrawal AA, Relyea RA. Re-evaluating the costs and limits of adaptive phenotypic plasticity. Proceedings of the Royal Society B: Biological Sciences. 2010;277(1681):503–511. doi:10.1098/rspb.2009.1355.

34. Murren CJ, Auld JR, Callahan H, Ghalambor CK, Handelsman CA, Heskel MA, et al. Constraints on the evolution of phenotypic plasticity: limits and costs of phenotype and plasticity. Heredity. 2015;115(July 2014):293–301. doi:10.1038/hdy.2015.8.

35. Kashtan N, Mayo AE, Kalisky T, Alon U. An Analytically Solvable Model for Rapid Evolution of Modular Structure. PLoS Computational Biology. 2009;5(4). doi:10.1371/journal.pcbi.1000355.

36. Kouvaris K, Clune J, Kounios L, Brede M, Watson RA. How evolution learns to generalise: Using the principles of learning theory to understand the evolution of developmental organisation. PLOS Computational Biology. 2017;13(4):e1005358. doi:10.1371/journal.pcbi.1005358.

37. Draghi JA, Whitlock MC. Phenotypic plasticity facilitates mutational variance, genetic variance, and evolvability along the major axis of environmental variation. Evolution. 2012;66(9):2891–2902. doi:10.1111/j.1558-5646.2012.01649.x.

38. Watson RA, Wagner GP, Pavlicev M, Weinreich DM, Mills R. The evolution of phenotypic correlations and “developmental memory”. Evolution. 2014;68(4):1124–1138. doi:10.1111/evo.12337.

39. West-Eberhard MJ. Developmental plasticity and evolution. Oxford University Press; 2003.

40. Scheiner SM, DeWitt TJ. Phenotypic plasticity: functional and conceptual approaches; 2004.

41. Hancock PJB. An empirical comparison of selection methods in evolutionary algorithms. In: AISB Workshop on Evolutionary Computing. Springer; 1994. p. 80–94.

42. Lipson H, Pollack JB. Automatic design and manufacture of robotic lifeforms. Nature. 2000;406(6799):974–978.

43. De Jong G, Bijma P. Selection and phenotypic plasticity in evolutionary biology and animal breeding. Livestock production science. 2002;78(3):195–214.

